# Seagrass beds provide habitat for crabs, shrimps and fish in two estuaries on the South Island of New Zealand

**DOI:** 10.1101/2020.07.22.120055

**Authors:** Mads S. Thomsen, Averill Moser, Micaela Pullen, Derek Gerber, Sarah Flanagan

## Abstract

Seagrasses are marine angiosperms that potentially provide habitat for crabs, shrimps and fish. However, these types of data are lacking for the seagrass species (*Zostera muelleri*/rimurēhia) that inhabit intertidal estuaries on the South Island of New Zealand.
Abundances of crabs, shrimps and fish were therefore quantified from 361 non-destructive seine tows done in seagrass beds and bare mudflats in Duvauchelle bay and two sites in the Avon-Heathcote/Ihutai estuary between October 2019 and February 2020.
A total of 2549 crabs, 5824 shrimps and 1149 fish (75% were juvenile flounders) were identified and counted in the seine-net and immediately released back in healthy condition to the exact location from where they were caught.
Only few seagrass leaves were caught in the net and these leaves may have been previously uprooted drift fragments. The instant catch-and-release methodology therefore leaves, literally, nothing but a footprint.
More fish taxa, including two species of pipefish, were found in seagrass beds in Duvauchelle bay than in the Avon-Heathcote estuary. Fish (minus juvenile flounders) were also more abundant in these seagrass beds. Furthermore, juvenile flounders and shrimps were more abundant in Duvauchelle bay compared to the Avon-Heathcote estuary, but were found in similar abundances in seagrass beds and on bare flats.
It is possible that more fish were found in Duvauchelle seagrass beds because these beds have adjacent deeper areas, and may have high connectivity to seagrass beds in nearby bays. This hypothesis should be tested by sampling more seagrass beds in different types of estuaries and bays.
By contrast, crabs were more abundant in the Avon-Heathcote estuary, where spider crabs were most abundant in the seagrass beds, but other crabs were found in similar abundances in seagrass beds and bare habitat. We hypothesize that crab abundances were higher in the Avon-Heathcote estuary because of lower fish predation pressure and/or larger populations of prey like mollusc and polychaetes.
Our results suggests that (a) superficially similar *Zostera* beds in relatively close proximity can provide very different habitat values for fish and crustaceans, (b) seagrass beds with higher diversity and abundances of fish may be prioritized in conservation and management (assuming other important ecosystem functions are similar between beds), and (c) that pipefish may be useful indicator organisms, representing healthy, extensive, dense and connected seagrass beds.

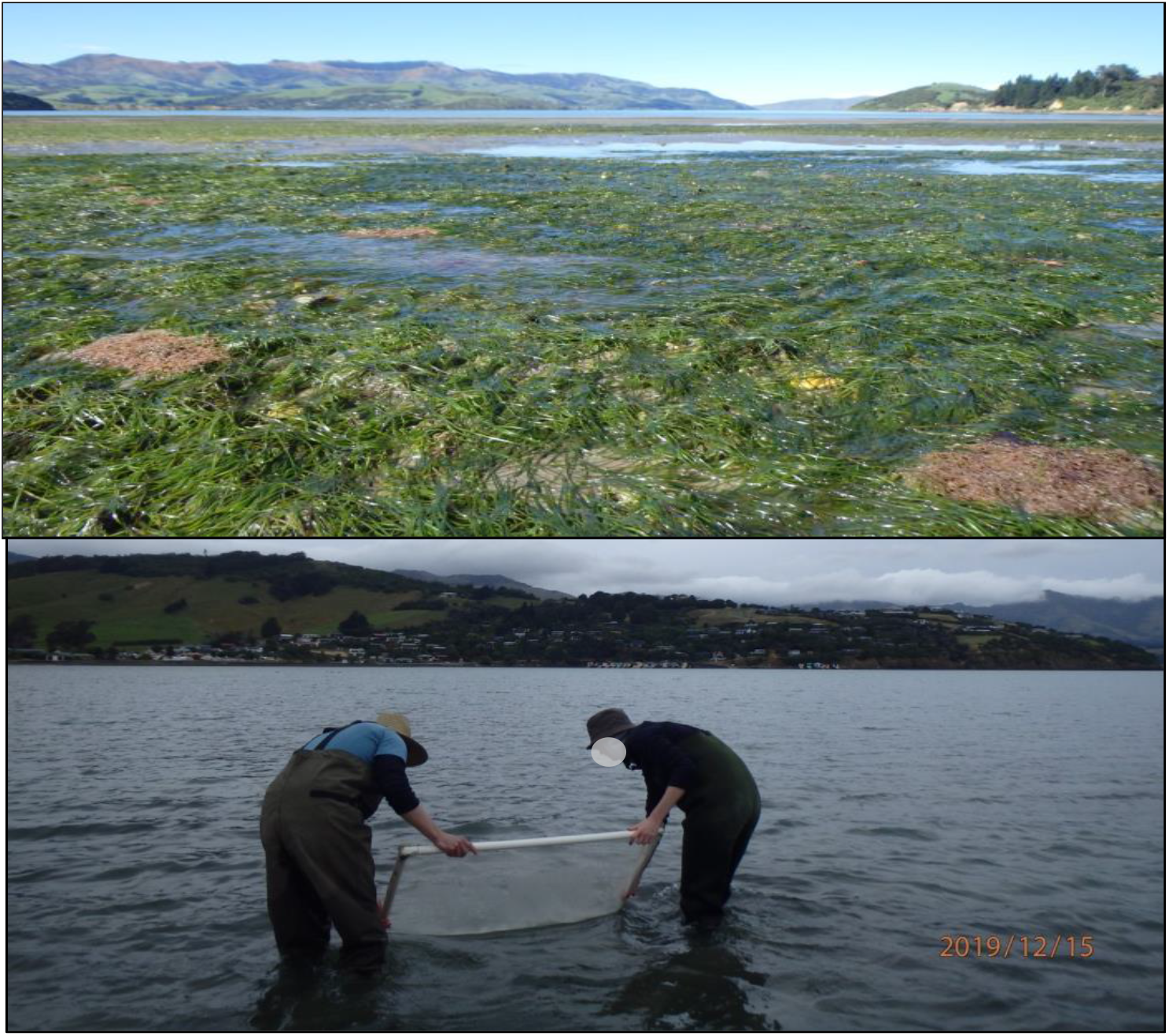

## Introduction

Seagrasses are marine angiosperms that provide important ecosystem functions and services (Hemminga and Duarte 2000, Cullen-Unsworth and Unsworth 2013, Nordlund et al. 2016, Nordlund et al. 2018). For example, seagrass can attenuate waves, stabilize sediments, reduce erosion, sequester carbon, take up land-derived nutrients and provide habitat for epiphytes, invertebrates and fish (Ward et al. 1984, Koch and Gust 1999, Holmer et al. 2004, Nayar et al. 2010, Thomsen et al. 2010, Tuya et al. 2011, Fourqurean et al. 2012, Thomsen et al. 2018). More specifically, seagrasses are considered important ‘nursery habitats’ for fish and crustaceans, where early life stages can find ample food and escape predation (Haywood et al. 1995, Heck et al. 2003, Lilley and Unsworth 2014). However, many ecosystem functions and services have been reduced or lost as seagrass around the world have been decimated due to human-associated activities, such as eutrophication, competition from nutrient-fuelled epiphytes and seaweeds, land-reclamation, habitat-alterations, global warming and heatwaves, and interactions with non-native species (Waycott et al. 2009, Hoeffle et al. 2011, Holmer et al. 2011, Hoeffle et al. 2012, Thomsen et al. 2012, Arias-Ortiz et al. 2018, Siciliano et al. 2019, Smale et al. 2019).

Seagrass beds have, for the same reasons, also been reduced throughout New Zealand, although the extents of losses is still relatively unknown (Inglis 2003, Turner and Schwarz 2006, Matheson et al. 2011). A singles seagrass species, *Zostera muelleri/*rimurēhia (hereafter *Zostera*), exists in New Zealand (Les et al. 2002), with distinct genetic populations in different parts of the country (Jones et al. 2008). Most of these populations occupy intertidal and upper subtidal mud and sand-flats in shallow estuaries (Inglis 2003, Turner and Schwarz 2006, Matheson et al. 2011), although deeper populations can also be found in more open sandy areas with high water clarity and reduced wave actions (Schwarz et al. 2006, Booth 2019). Relatively few studies have investigated whether *Zostera* in New Zealand provide habitat for crabs, shrimps and fish compared to adjacent mudflats, and most of these studies have been done around subtidal seagrass beds on the North Island (Parsons et al. 2014, Cooper 2015, Parsons et al. 2015, Parsons et al. 2016)-but see (Morrison et al. 2014) for a rare example from the South Island. The objective of this report is to address this research gap and quantify with seine tows (Fig. 1) if seagrasses in Duvauchelle bay and the Avon-Heathcote estuary/Ihutai, on the South Island of New Zealand, provide habitat for crabs, shrimps and fish, and if so, if abundances are different from adjacent mudflats.

**Fig. 1.**
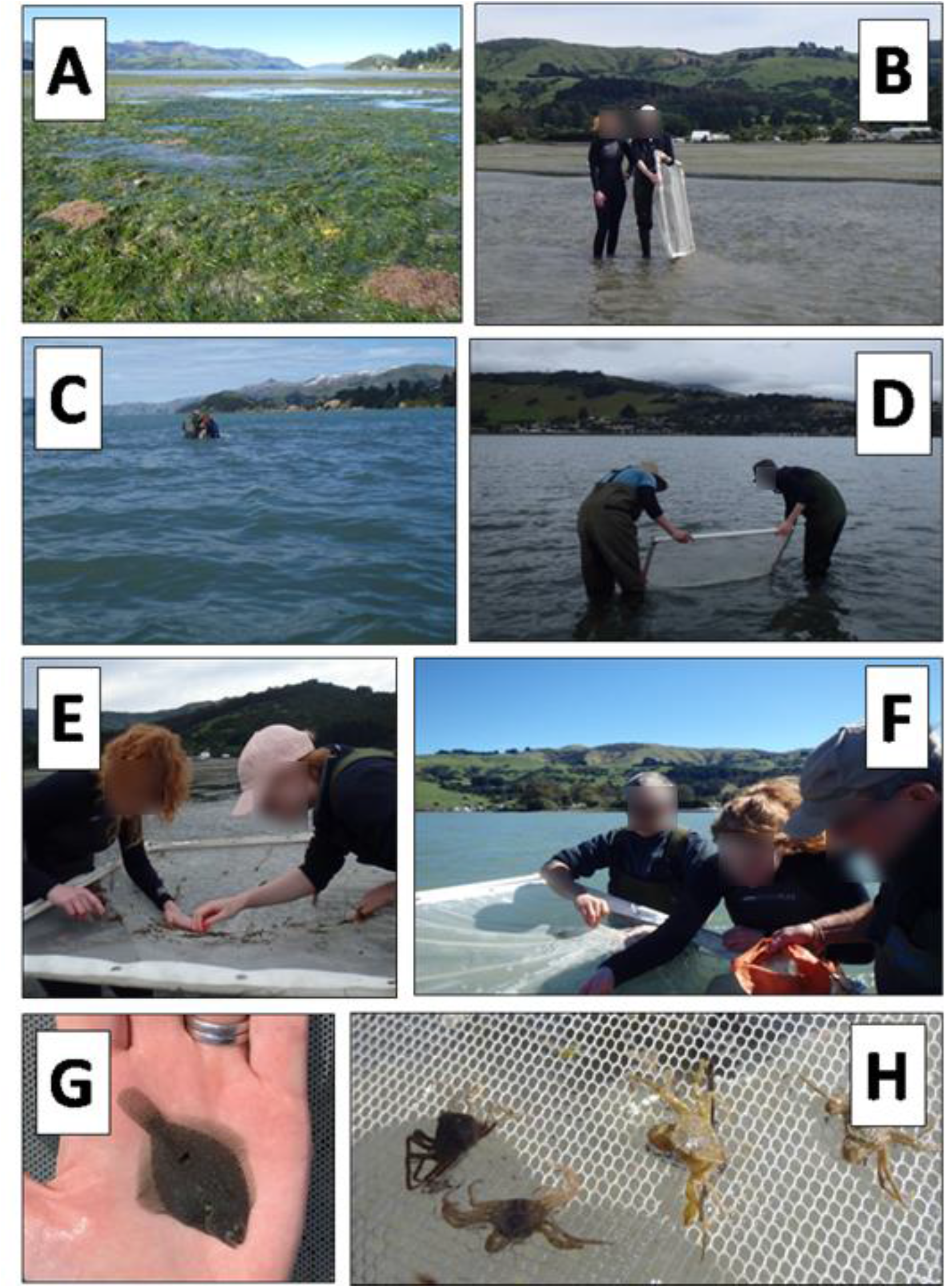
Photos showing (A) typical seagrass bed in Duvauchelle bay, (B) the small and flexible seine net, (C) seining in deep and (D) shallow waters, (E, F) identification, counting and release of organisms at the collection site, and examples of common organisms, including (G) juvenile flounder and (H) *Halicarcinus* spp..

## Methods

### Study sites

The Avon-Heathcote estuary is a ca. 8.8 km^2^ shallow, well-flushed, bar-built estuary surrounded by the city of Christchurch (Fig. 2). Two rivers flow into the estuary; the Avon River flows from the north and the Heathcote from the southwest, creating a salinity gradient ranging from ca. 8-15 psu around the mouth of the rivers to c. 22-34 psu near New Brighton Spit and the estuary mouth (Marsden 2004). Seagrass beds are abundant along the eastern side of the estuary, covering ca. 0.35 km^2^ (Hollever and Bolton-Ritchie 2016). The tides ranges from 1.7 m at neap to 2.2 m at spring tide (Knox et al. 1973).

**Fig. 2.**
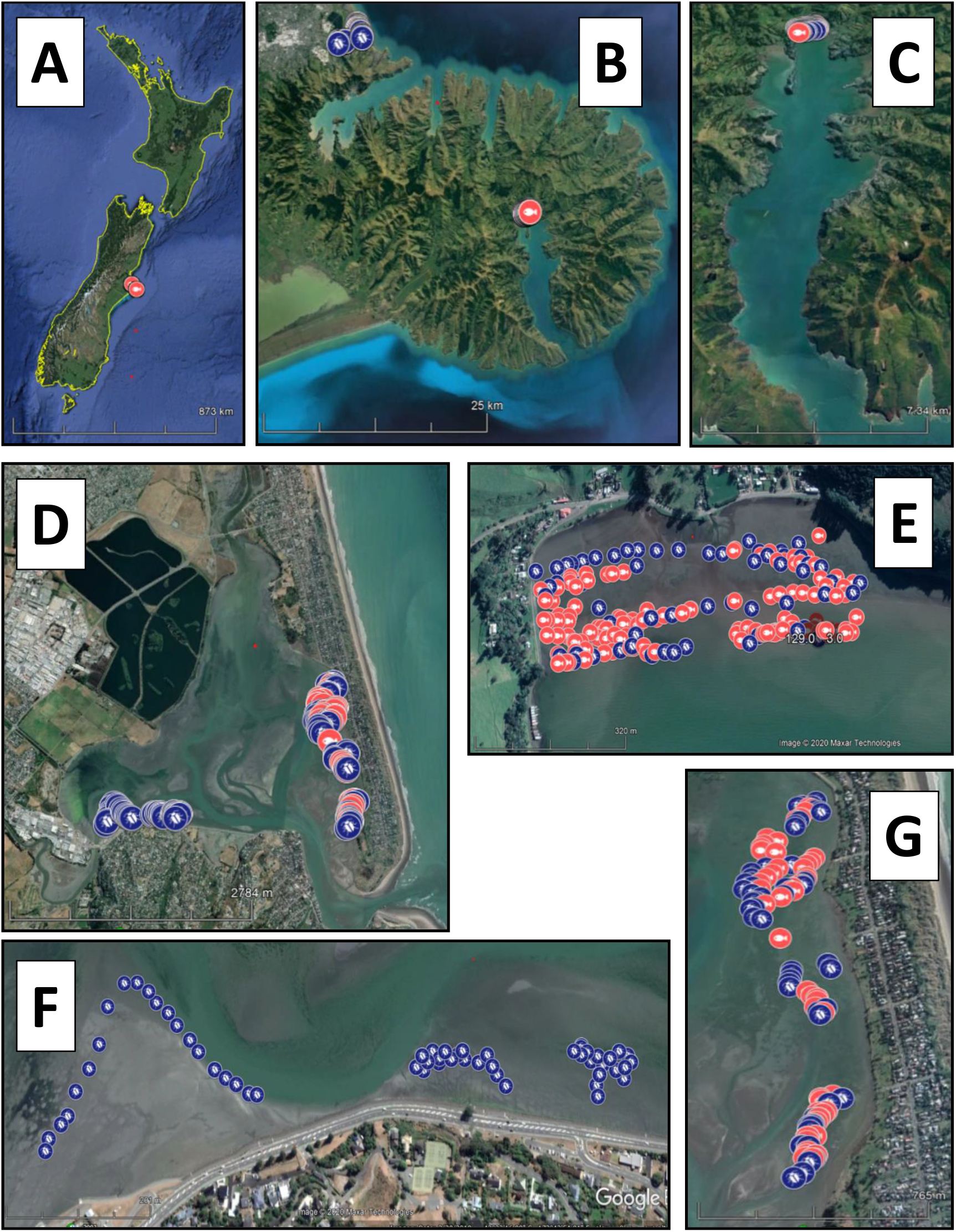
Spatial position of seine tows in seagrass beds (red) and bare sediment flats (blue) across spatial scales ranging from New Zealand (A), Banks Peninsula and Avon-Heathcote estuary (B), Akaroa Harbour within Banks Peninsula (C), Avon-Heathcote estuary (D), Duvauchelle bay within Akaroa Harbour (E) and the Causeway (F) and Spit (G) within the Avon-Heathcote estuary. See Fig. 3 for number of tows per habitat.

Duvauchelle bay is the northernmost, furthest inland bay in Akaroa Harbour, situated approximately 16 km from the open ocean (Fig. 2). The bay is 300 m wide and in the nearshore area it is very flat, descending 1.5 m over 800 m (Hart et al. 2009). This location makes it one of the most protected bays in Banks Peninsula, and it is a large mudflat dominated by sands (Hart et al. 2009) with seagrass beds throughout the intertidal zone. A stream flows into the bay about halfway through the bay, creating a natural barrier between the two halves of the bay (Fig. 2). The upper Akaroa Harbour intertidal mudflats can be considered ‘Areas of Significant Natural Value’ (Bolton-Ritchie 2005).

### Seine-sampling

Sampling was done with a modified 1.2 × 1.2 m seine net mounted on a PVC frame with a mesh size of 0.5 mm (Fig. 1). The net was hand-hauled, parallel to the shore with a slope of approximately 60-80 degrees. At the start and end of each of tow, GPS coordinates and the exact time were recorded on a Garmin GPS. The PVC frame allows the net to be balanced between two people while in the water. A total of 361 tows were done in seagrass beds and bare mudflats in Duvauchelle bay and two sites in the Avon-Heathcote estuary between October 2019 and February 2020 (see Fig. 2 and 3 for details of locations and number of tows from different sites and habitats). Water depth varied from 0.15-1.2 m and towing length and duration typically ranged between 3-15 m and 1-15 minutes depending on local conditions, that is, some tows were short (and therefore fast) to remain within a small sand or seagrass patch. Sampling was done between 2 hours before low tide and 5 hours after low tide. After each tow, we counted and identified fish, crabs and shrimps to operational taxonomic units, directly on the seine-net at the GPS location of capture. Organisms were immediately released alive and healthy after identification and counting to minimize environmental impacts. This instantaneous and non-destructive ‘on-the net’ sampling meant that many small and juvenile organisms could only be identified to taxonomic units above the species levels. We also recorded if live seagrass shoots were caught in the net to estimate it the net disturbed the bed by uprooting leaves.

**Fig. 3.**
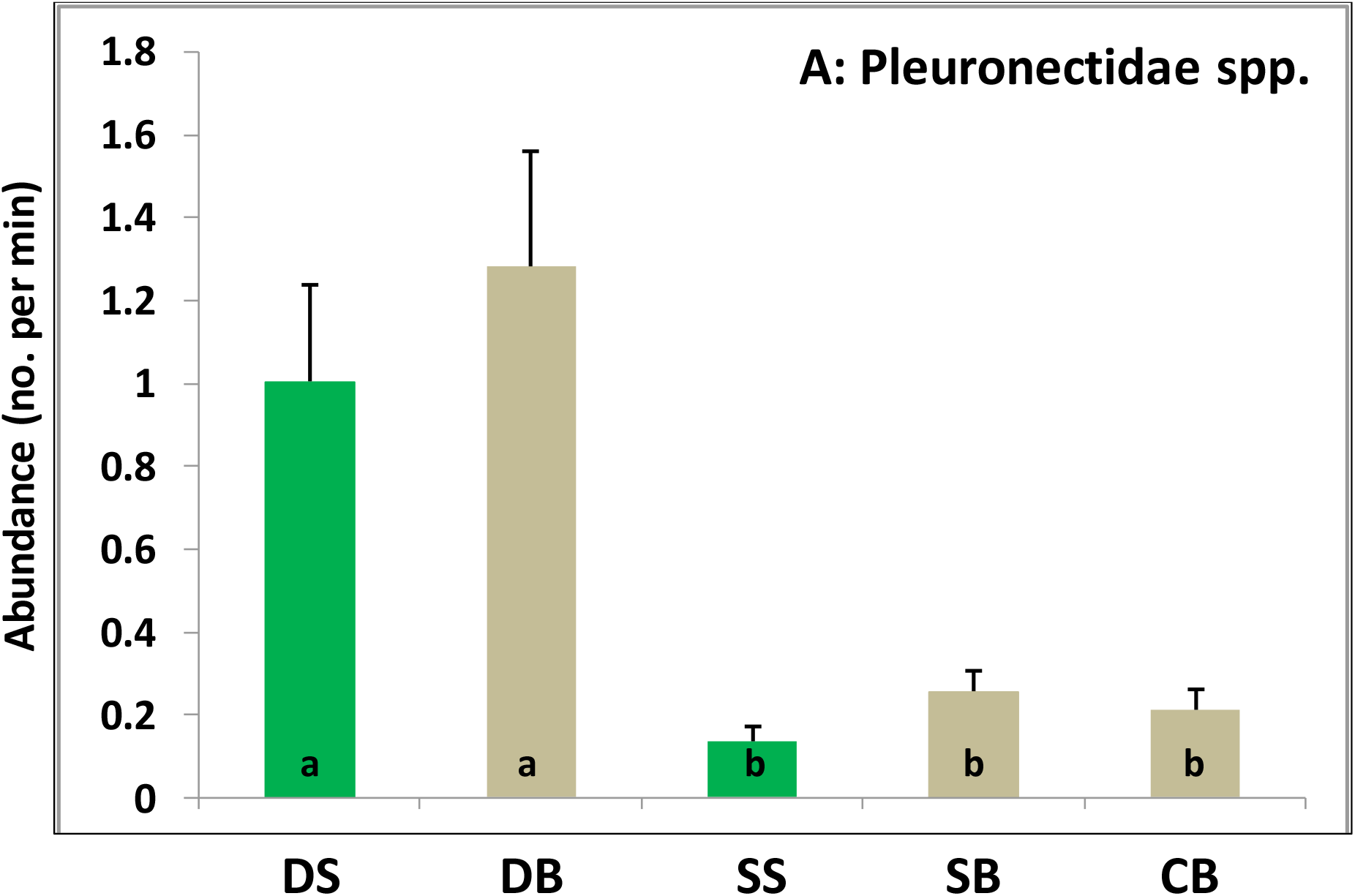
Abundances of flounders (+ SE) caught in seine tows (number per minute of towing). Lower capital letters on bars = significant differences between five sampled habitats; DS = Duvauchelle bay seagrass bed (n = 129), DB = Duvauchelle bay bare flat (n = 63), SS = Spit seagrass bed (n = 55), SB =Spit bare flat (n = 50), and CB = Causeway bare flat (n = 60). See figure 2 for map and locations of seine tows.

### Statistical analysis

. All abundances per tow were standardized to unit time, i.e., numbers of organisms per minute of towing (this is a more precise estimate of sampling effort compared to GPS-calculated distances that have positional errors associated with both the start and stop locations). We then tested if standardized abundances of fish, shrimps and crab taxa differed between five combinations of habitats (seagrass or bare flat) and sites (causeway and spit in the Avon-Heathcote estuary and Duvauchelle bay) with non-parametric Kruskal-Wallis tests. Non-parametric tests were used because data were non-normal, had many zeroes, and variances could not be transformed to homogeneity. Significant results were followed by Dunn’s post hoc comparisons. Significance was evaluated at alpha = 0.05.

## Results

A total of 2549 crabs, 5824 shrimps and 1149 fish were caught in the 361 seine tows. Flounders were, by far, the most abundant fish (75% of all fish) and almost all these flounders were juveniles that were smaller than 10 cm. There were significant differences between the five site/habitats (p = 0.01) with more fish in Duvauchelle bay than the Avon-Heathcote estuary, but with no differences between seagrass and bare habitats (Fig. 3). Most other fish taxa, including *Leptonotus elevatus* (high-bodied pipefish, p < 0.01, Fig. 4A), *Stigmatopora nigra* (wide-bodied pipefish, p < 0.01, Fig. 4B), blenniformes spp. (p < 0.01, Fig. 4C), *Arripis trutta* (kahawai, p = 0.02, Fig. 4D) and *Galaxias* spp. (‘whitebait’, p = 0.05, Fig. 4E), were significantly more abundant in seagrass beds in Duvauchelle bay compared to other habitats and sites. A few *Notolabrus celidotus* (spotty) individuals were found only in the seagrass beds of Duvauchelle bay, but this result was not significant (p = 0.46, Fig. 5F). In short, the two pipefish species, blenniformies spp., *A. trutta* and *N. celidotus* were only found in Duvauchelle bay and not the Avon-Heathcote estuary.

**Fig. 4.**
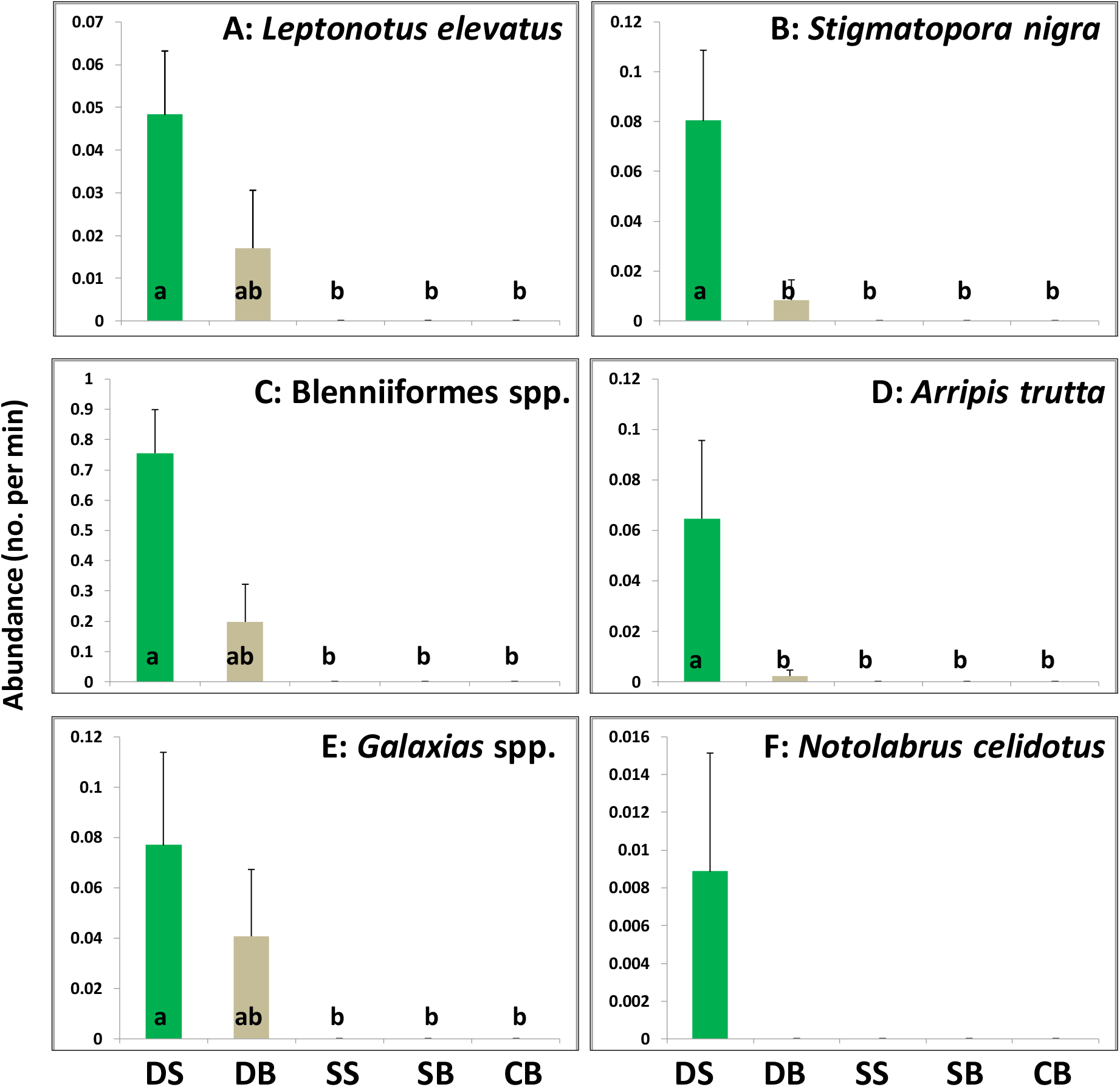
Abundances of fish (minus flounders, + SE) caught in seine tows (number per minute of towing). Lower capital letters on bars = significant differences between five sampled habitats; DS = Duvauchelle bay seagrass bed (n = 129), DB = Duvauchelle bay bare flat (n = 63), SS = Spit seagrass bed (n = 55), SB =Spit bare flat (n = 50), and CB = Causeway bare flat (n = 60). See figure 2 for map and locations of seine tows.

**Fig. 5.**
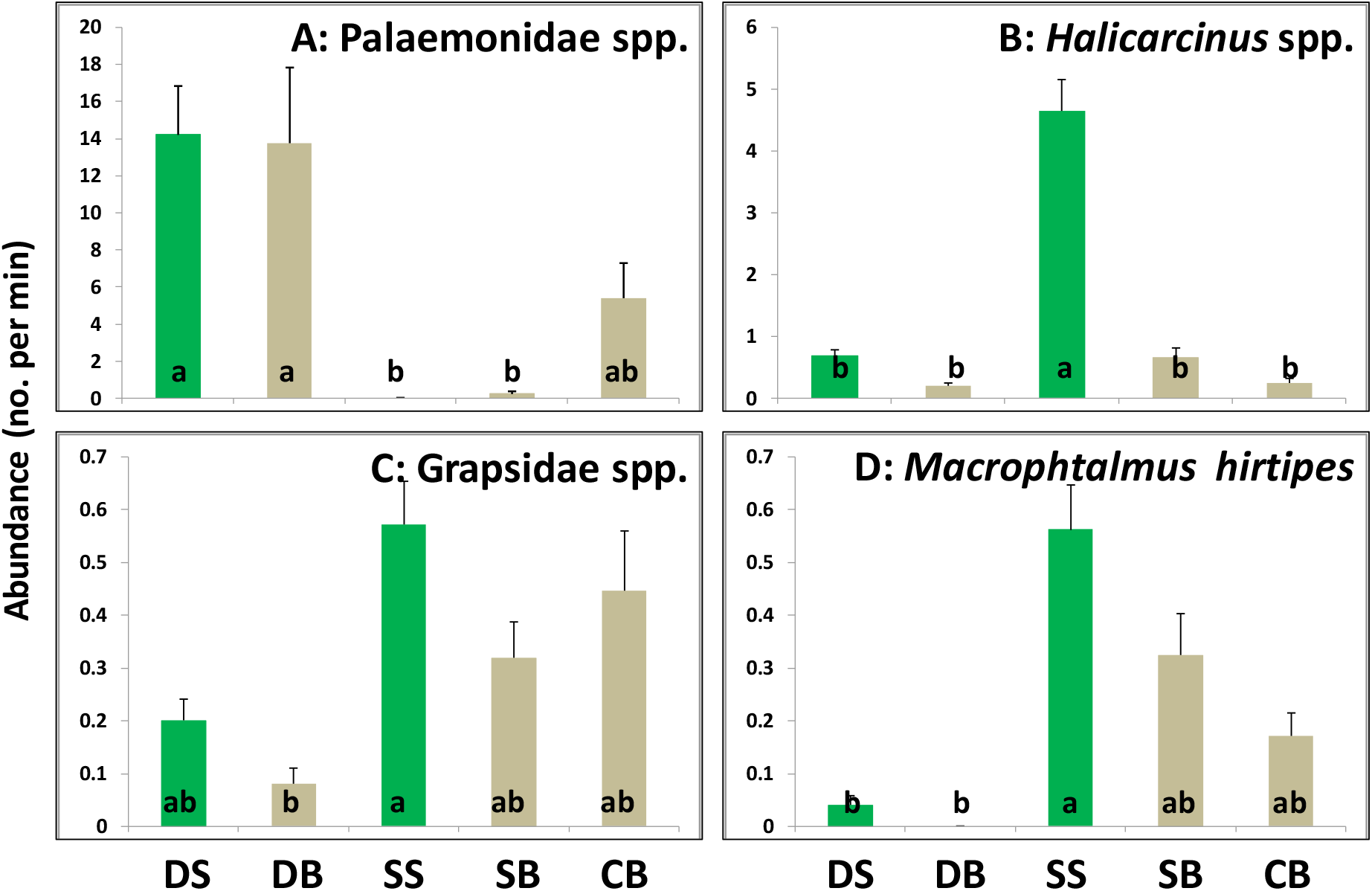
Abundances of crabs and shrimps (% SE) caught in seine tows (number per minute of towing). Lower capital letters on bars = significant differences between five sampled habitats; DS = Duvauchelle bay seagrass bed (n = 129), DB = Duvauchelle bay bare flat (n = 63), SS = Spit seagrass bed (n = 55), SB =Spit bare flat (n = 50), and CB = Causeway bare flat (n = 60). See figure 2 for map and locations of seine tows.

All invertebrate abundances were significant (p < 0.01); *Palaemonidae* spp. (shrimps) were most abundant in Duvauchelle bay (but with no differences between habitats, Fig. 5A), *Halicarcinus* spp. (spidercrabs) was most abundant in the seagrass beds on ‘the spit’ in the Avon-Heathcote estuary (Fig. 5B), and grapsid crabs (various mud crabs, Fig. 5C) and *Macrophtalmus hirtipes* (stalk-eyed mud crab, Fig. 5D) were most abundant in the Avon-Heathcote estuary (but, again, with no differences between habitats).

## Discussion

### Discussion of results

We found striking differences in fish and crab populations between two estuaries situated in relatively close proximity (Fig. 2) experiencing similar external climatic and oceanographic conditions. However, analogue differences in seagrass nekton communities within biogeographical regions have been observed before in both Northern and Southern New Zealand (Morrison et al. 2014). We generally found few fish taxa compared to Morrison et al.’s fish survey, in part because they used longer tows with a much larger net (11 m wide × 2.3 m high, each tow covering ca. 450 m^2^ of seafloor), in part because northern estuaries generally have much more diverse and abundant fish populations (e.g., >2000 juvenile snappers have been caught in a single tow in Raglan harbour) (Morrison et al. 2014).

Our results support past studies from around the world (Heck et al. 2003, Lilley and Unsworth 2014, Parsons et al. 2015) and New Zealand (Lowe 2013, Morrison et al. 2014) in that fish populations generally were larger and more diverse in seagrass beds compared to adjacent bare flats, probably because predation pressure are lower and food levels higher in the seagrass bed. However, this pattern was only clear in Duvauchelle bay, i.e., the seagrass beds in the Avon-Heathcote were inhabited by very few fish (minus flounders). This is unlikely to be attributed to small scale seagrass attributes because leaf length, width and densities, and seagrass patch sizes, are relatively similar between the two estuaries (Thomsen, unpublished data). The striking difference is more likely related to larger scale attributes. For example, diversity and abundance of nekton can vary with seagrass landscapes characteristics, including patch size, shape, and configuration, landscape connectivity, and estuarine geomorphology, hydrology, catchment characteristics, and human land-use and fishing patterns (Brooks and Bell 2001, Hovel and Lipcius 2001, Salita et al. 2003, Hovel and Fonseca 2005, Boström et al. 2006a, Boström et al. 2006b, Connolly and Hindell 2006, Jelbart et al. 2006, Gullström et al. 2008, Boström et al. 2010, Tuya et al. 2010, Smith et al. 2011b).

We found several native and endemic species only in Duvauchelle bay, including two pipefish species. Unlike many fish that use seagrass habitats as a nursery ground, pipefish species such as *S. nigra* spend their entire lives amongst the seagrass (Bell and Westoby 1986, Steffe et al. 1989, Browne 2008, Smith et al. 2011a), relying on the seagrass for shelter from predators and strong wave action (Hindell et al. 2000a, Hindell et al. 2000b, 2001, Moran et al. 2003, Smith et al. 2010, Smith et al. 2011a, Smith et al. 2011b). In Australia, *S. nigra* are known to prefer dense, tall seagrass and are one of the most abundant fish in the seagrass meadows (Edgar and Shaw 1995, Hindell et al. 2000b, Macreadie et al. 2009, Smith et al. 2010). Given these characteristics of this family of fishes (Syngnathidae), and the fact that they are easy to sample with a small net such as the one used here, we recommend that pipefish be used as indicator organisms for management and conservation in New Zealand. The presence of pipefish would suggest that the seagrass meadow is healthy and well-connected, and would be indicative of high species richness (Shokri et al. 2009). This suggest that the Avon-Heathcote contains lower-quality seagrass beds from an ecological perspective, possible reflecting differences in water quality (e.g., nutrient load) and high abundances of drift seaweed (Bolton-Ritchie 2005, Bolton-Ritchie and Main 2005, Bolton-Ritchie 2008) and/or differences in large scale attributes such as catchment usage, geomorphology and connectivity.

Shrimp and juvenile flounder dominated the seagrass community in Duvauchelle bay but not in the Avon-Heathcote estuary. High abundances of juvenile flounder have been recorded in other estuaries in New Zealand, particularly in areas with soft sediments (Lowe 2013), which are characteristic of Duvauchelle bay. Both juvenile flounder and pipefish prefer mysids as prey (Smith et al. 2011a, Lowe 2013), so it is possible that differing zooplankton communities explain the differences we observed between estuaries. Shrimps generally select habitats based on a variety of visual and chemical cues (Lecchini et al. 2010), and features such as habitat complexity and predator presence are known to determine shrimp microhabitat selection (Tait and Hovel 2012). Therefore, it is likely that the differences we observed are due to variation in unmeasured habitat characteristics between Duvauchelle bay and the Avon-Heathcote estuary. Future work could investigate whether disparities in the communities correspond to differences in factors such as turbidity, zooplankton communities, and variability in environmental factors.

The result for the benthic crabs differed from the pattern of the nektonic fish and shrimps, because crabs were generally more abundant in the Avon-Heathcote estuary, perhaps because there are less predation by nektonic fish (as suggested but the present study) and/or more food (molluscs and worms are more common in the Avon-Heathcote estuary (Siciliano 2018, Foster 2019). Of the different crab taxa *Halicarcinus* was particularly abundant in the seagrass bed in Avon-Heathcote estuary. These small and cryptic herbivorous crabs are susceptible to predation because of their small chelae, slow movement and soft exoskeleton, and are therefore likely to rely on seagrass (with few fish) for hiding and feeding on small epiphytes on the seagrass leaves (Bell et al. 1978, Klumpp and Nichols 1983, van Houte-Howes et al. 2004, Vinuesa et al. 2011).

### Implications for conservation

Our results highlight that superficially similar *Zostera* beds in relatively close proximity can provide very different habitat values for fish and crustaceans. Therefore, prioritizing the conservation and management of seagrass beds with higher diversity and abundances of fish will benefit a larger number of native species, assuming other important ecosystem functions are similar between beds. We further recommend that pipefish species can be used as indicators of the quality of the ecosystem function of seagrass habitats in New Zealand.

### Study limitations, environmental impact and recommendations for future studies

Our sampling methodology had several limitations that can bias the results. First, there are well-established biases associated with the net seize, type, mesh-size, frame, towing speed, towing angle, etc. (Penczak and O'Hara 1983, Pierce et al. 1990, Weaver et al. 1993, Guest et al. 2003, Franco et al. 2012). For example, our seine net is ‘small’ compared to nets typically used in fish-surveys, implying that fish may escape to the side of the net. Furthermore, because the seine is hand-towed, it can only be operated to a maximum of ca. 1.5 m depth, limiting the depth of sampling compared to boat trawling. Because of the small net size and slow hand-towing, highly mobile and larger species are more likely to escape and thereby be under-sampled. Finally, it is possible that some crabs and fish can cling tightly to leaves that bend under the seine and thereby escape detection.

However, there are also advantages of our methodology, including instantaneous identification and counting of organisms on the net at the exact location of capture. For example, the small net size makes the net easy and flexible to transport, carry and operate by two persons (with a third person to write) allowing for sampling in difficult to reach areas in both small and large patches. The small mesh size also ensures that juvenile organisms are sampled. This method is therefore particularly useful in shallow wave-protected areas (like estuaries) and for targeting organisms that tend to ‘hide’ rather than ‘flee’ (like many seagrass inhabiting taxa) (Penczak and O'Hara 1983, Pierce et al. 1990, Guest et al. 2003). Thus, we expect to have more reliable estimates of organisms like pipefish, shrimps and arrow crabs compared to (underestimated) juvenile flounders and wrasses. Another major advantage of our methodology is its small environmental impact. The captured organisms typically remained submerged during counting and were released back to the exact location of capture. We observed movement patterns and behaviours of release organisms, and they appeared unharmed with no visible damage and were observed swimming and crawling back to their habitat. Furthermore, we recorded few live seagrass leaves on the net, implying that the seagrass bed was not affected by the disturbance. Indeed, because we sometimes observed leaves in bare mudflat tows it is possible that leaves primarily were drift-fragments. Indeed, drifting leaves are common in estuaries and on many beaches and represent an important dispersal mechanism (Jones et al. 2008, Berković et al. 2014, Stafford-Bell et al. 2015). Captured leaves were also released so that they, if they had roots and rhizomes, could re-attached into the sediment or spread to nearby areas (Ewanchuk and Williams 1996, Berković et al. 2014, Stafford-Bell et al. 2015). Thus, the main impact of out seining is the ‘footprint’ of the samplers, but we have generally observed, from years of studying seagrasses, that *Zostera muelleri* recover rapidly from a foot print (pers. obs.), although intensive and repeated trampling associated with fishing and digging obviously have stronger and more lasting impacts (Eckrich and Holmquist 2000, Alexandre et al. 2005, Skilleter et al. 2006, Travaille et al. 2015, Garmendia et al. 2017).

In addition to biases associated with the sampling methodology, there are also biases associated with our spatio-temporal sampling design. For example, we did not take into account the exact water depth, time of day, time related to the tidal flow, seasonality, or seagrass traits such as leave density, length and width, patch size and patch configuration, presence of epiphytes or drift seaweed, etc., attributes that can affect abundances of decapod and fish nekton (Haywood et al. 1995, Heck et al. 2003, Jelbart et al. 2006, Smith et al. 2010, Smith et al. 2011a, Lilley and Unsworth 2014, Thomsen et al. 2018). Based on the limitations outlined above, we recommend additional seining to better understand the distributions of nektonic crustaceans and fish, to allow managers to priorities conservation of specific seagrass beds and habitats and to aid future mitigations and restoration efforts. For example, future seining should including sampling (i) over shorter and longer time scales (such as day-night, morning-evening, lunar cycles, seasons, and across years), (ii) across small scale habitat gradients (e.g., gradient in seagrass densities, leave lengths, patch sizes, drift alga, or degree of epiphytism), (iii) medium scale spatial gradients (within estuaries such as along depth, salinity, turbidity and sediment gradients), and (iv) along large scale spatial gradients (between estuaries, such as across latitudes, sizes and types of catchment areas, geomorphology and tidal flow regimes, and degree of adjacency and connectivity to other estuaries and seagrass beds). Finally, future seining should also sample nekton that have experienced anthropogenic-associated stressors, such as fishing, drift weed accumulations, eutrophication, enhanced sediment loadings, and marine heatwaves, to better understand human impacts and improve future management and conservation of seagrass beds.

## Conclusion

Juvenile flounders were, by far, the most common fish taxa, being particularly abundant in Duvauchelle bay, in both seagrass and bare habitats. We found the highest biodiversity and abundances of other fish taxa, including two pipefish species, in the seagrass beds of Duvauchelle bay. Furthermore, shrimps were also most abundant in Duvauchelle bay, whereas spider crabs were more abundant in the seagrass beds in the Avon-Heathcote estuary. Finally, we found that other crabs were most abundant in the Avon-Heathcote estuary but with similar abundances in seagrasses and bare habitats. We hypothesize that more fish taxa were found in the seagrass beds in Duvauchelle bay because these beds have adjacent deeper areas and may be better connected by migrations to seagrass beds in nearby bays. These results highlight that *Zostera* beds within the same biogeographical region can provide very different habitat values. We finally suggest that similar surveys are done in estuaries throughout New Zealand to identify seagrass beds of highest conservation priorities, to aid management of contemporary seagrass beds and to monitor future impacts from anthropogenic stressors.

## Acknowledgement and permits

This research was supported by a University of Canterbury Summer Scholarship sponsored by Environment Canterbury to Averill Moser. Sampling was conducted in accordance to MPI Special Permit 590 and Special Permit Amendment 590-2 to the University of Canterbury.

## Notes

### Competing Interest Statement

The authors have declared no competing interest.

